# Identifying complex motifs in massive omics data with a variable-convolutional layer in deep neural network

**DOI:** 10.1101/508242

**Authors:** Jing-Yi Li, Shen Jin, Xin-Ming Tu, Yang Ding, Ge Gao

## Abstract

Motif identification is among the most common and essential computational tasks for bioinformatics and genomics. Here we proposed a novel convolutional layer for deep neural network, named Variable Convolutional (vConv) layer, for effective motif identification in high-throughput omics data by learning kernel length from data adaptively. Empirical evaluations on DNA-protein binding and DNase footprinting cases well demonstrated that vConv-based networks have superior performance to their convolutional counterparts regardless of model complexity. Meanwhile, vConv could be readily integrated into multi-layer neural networks as an “in-place replacement” of canonical convolutional layer. All source codes are freely available on GitHub for academic usage.

## INTRODUCTION

Recurring sequence motifs [1, 2] have been well demonstrated to exert or regulate important biological functions, such as protein binding [3], transcription initiation [4], alternative splicing [5], subcellular localization [6], translation control [7], and microRNA targeting [8]. Effectively and efficiently identifying these motifs in massive omics data is a critical step for follow-up investigations.

Various computational tools have been developed to identify sequence motifs via word-based and profile-based models [9-14]. Word-based tools start with a fixed-length and conservative segment and then perform a global scanning; such tools include DREME [15], Fmotif [16], RSAT peak-motifs [17], SIOMICS [18, 19], and Discover [20]. While these tools can theoretically obtain the globally optimal solution, they suffer from high computational complexity when applied to data with complex motifs or large-scale datasets [10]. Profile-based tools attempt to find representative motifs by heuristically fine-tuning a series of possible motifs, either generated from a subset of input data or randomly chosen [21-25], leading to a faster (but probably sub-optimal) motif calling [12].

Several convolutional neural network (CNN)-based tools have been proposed recently as a more scalable approach for identifying motifs. Alipanahi et al. developed DeepBind to identify protein binding motifs from large-scale ChIP-Seq datasets [26] by treating each convolutional kernel as an individual motif scanner and discriminating motif-containing sequences from others based on the output of all kernels. Along with this line, several convolution-based networks have been proposed to effectively handle large amount of data in various settings [27-34]. Meanwhile, however, the inherent fixed-kernel design of canonical convolutional networks could hinder effective identification [35-37] of *bona fide* sequence patterns, which are usually of various lengths and unknown *a priori* and often function combinatorially [38, 39]. One possible workaround is to build these representations in a hierarchical manner [28, 32], yet the deeper structure itself would demand additional time/space cost for network training and hyper-parameter search (also see Supplementary Notes 1).

Here, we proposed a novel convolutional layer for deep neural network called Variable Convolutional neural layer (vConv), which learns the kernel length directly from the data. Evaluations based on both simulations and real-world datasets showed that vConv-based networks outperformed canonical convolution-based networks, making it an ideal option for the *ab initio* discovery of motifs from high-throughput datasets. Further inspection also suggested the vConv layer being robust for various hyper-parameter setups. All source codes are publicly available on GitHub (https://github.com/gao-lab/vConv).

## MATERIAL AND METHODS

### Design and implementation of vConv

Without loss of generality, we focus on nucleotide sequences where each nucleotide can take only one of four alphabets (A, C, G, and T (for DNA) or U (for RNA)). vConv (Fig. 1) is designed to be a convolutional layer equipped with a trainable “mask” to adaptively tune the effective length of the kernel during training.

**Fig. 1.**
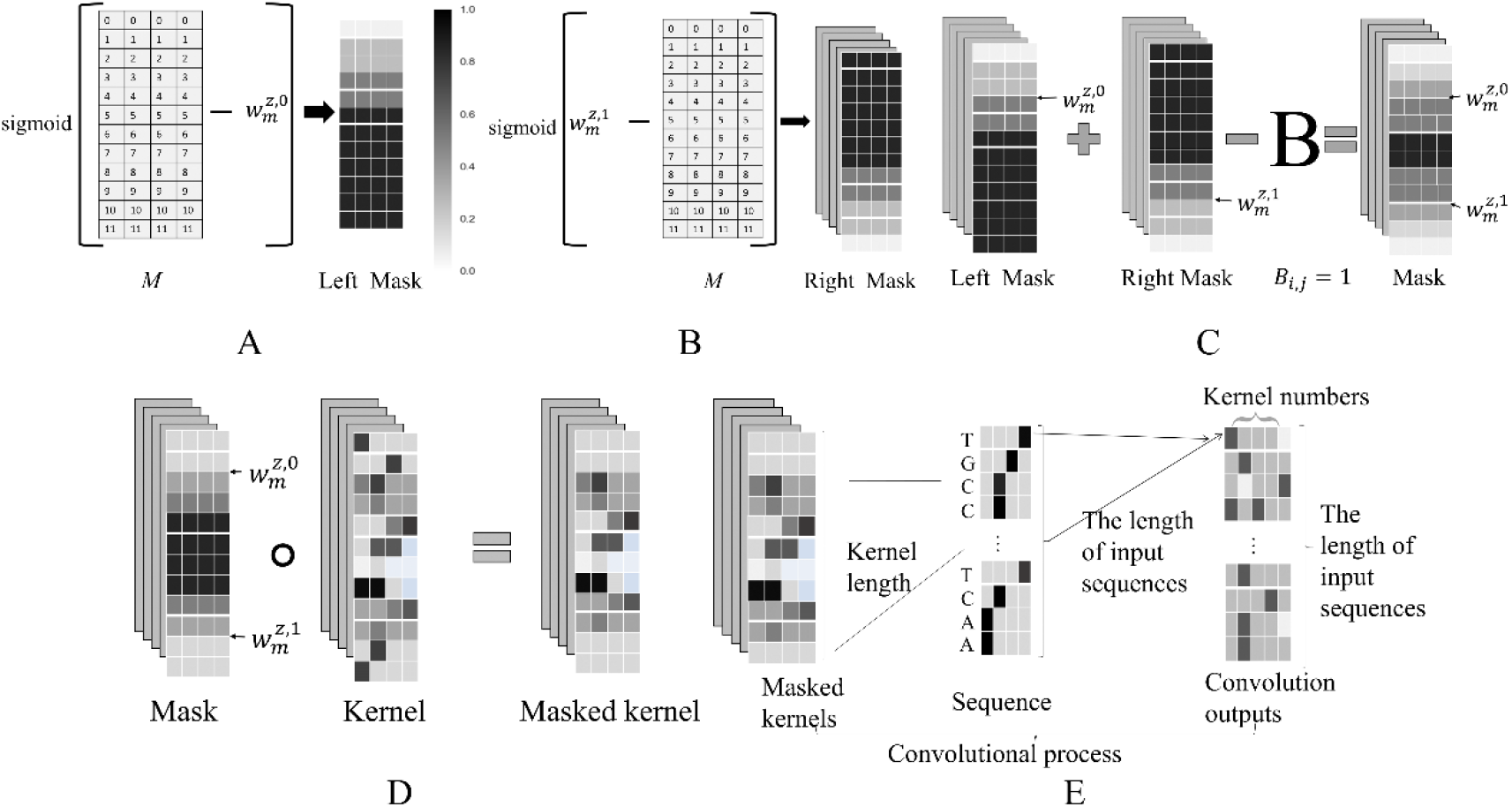
The design of vConv. To build up the mask matrix, vConv generates two sigmoid functions of opposite orientations (A and B), followed by overlaying them (C). Then vConv “masks” this mask matrix onto the kernel using the Hadamard product (D), and treats the masked kernel as an ordinary kernel to convolve the input sequence (E).

In brief, vConv uses the trainable left and right boundary values (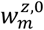 and 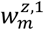, respectively) to generate two sigmoid functions 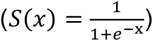 of opposite orientations (Fig. 1A-B), overlays them to define the final mask (Fig. 1C and Equation 5), uses this mask to mask a raw kernel (Fig. 1D), and finally uses the masked kernel to convolve sequences (Fig. 1E).

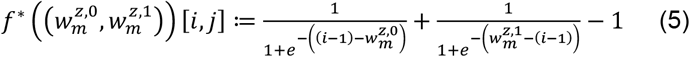

In practice, for a kernel with length *L*, the mask is generated as follows:

1. We first generate an *L* × 4 matrix *M* where *M*[*i, j*] = *i* − 1 (the grey numbered matrix in Fig. 1A-B);
2. We generate the first sigmoid 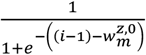 in Equation (5)) by subtracting 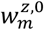 from *M* followed by a sigmoid transformation, thus pushing all those 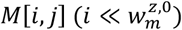 towards 0 and all those 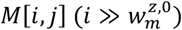 towards 1 (white and black cells in Fig. 1A, respectively);
3. Similarly, we generate the second sigmoid 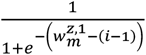 in Equation (5)) by adding 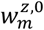 to −1 × *M* followed by a sigmoid transformation, thus pushing all those 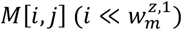 towards 1 and all those 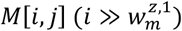 towards 0 (black and white cells in Fig. 1B, respectively);
4. Finally, we sum up the two sigmoid functions and shift all elements by -1 to restrict them within [0, 1] (Fig. 1C) to get the final mask.

One can multiply this mask and the original kernel by the Hadamard product ∘ (i.e.,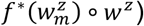) to effectively mask the raw kernel with the corresponding close-to-0 elements and obtain the masked kernel (Fig. 1D). This masked kernel can then be used as an ordinary kernel in a canonical convolutional layer (Fig. 1E). During training, the masking is essentially an operator with trainable weights (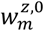 and 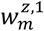) and will be automatically trained together with other weights.

As Equation 5 implies, all *i*’s falling outside the boundaries (i.e., 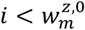 or 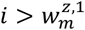) have their (masking) elements 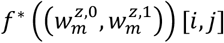 close to zero, and all *i*’s within the boundaries have their elements close to 1; those *i*’s around the boundaries have their elements around 0.5, thus ‘soft’-masking boundary kernel elements.

To make 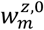 and 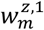 converge faster during training, we combine the binary cross-entropy (BCE) loss with a sum of masked Shannon losses (MSLs) from each kernel mask. Mathematically, we have the following total loss *L*:

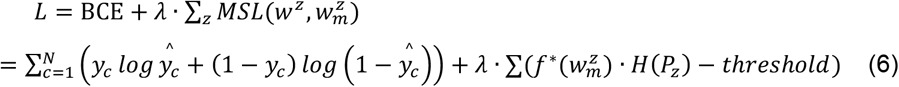

where 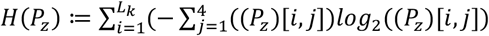 is the sum of the Shannon entropy across all nucleotide positions of *P*_*z*_, the position weight matrix (PWM) learned by the *z*-th kernel. For preciseness, we set *P*_z_ = *P*(*w*^*z*^, *b* = 2) , where 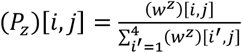 following the exact kernel-to-PWM transformation specified by Ding et al. [40]. One can then immediately deduce the formula for updating the left boundary 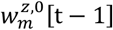 at time step *t* via gradient descent with learning rate *r* (*r*>0):

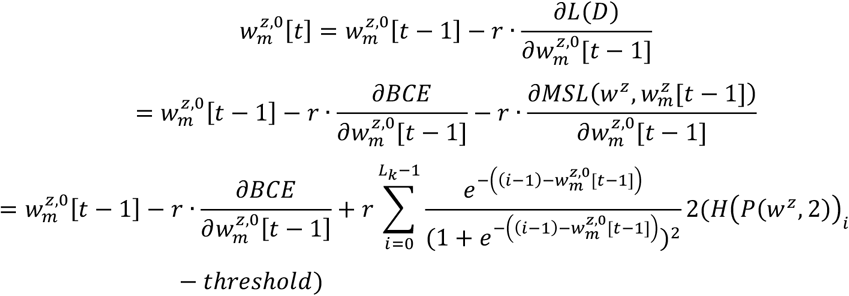

Similarly, for updating the right boundary 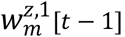, we have:

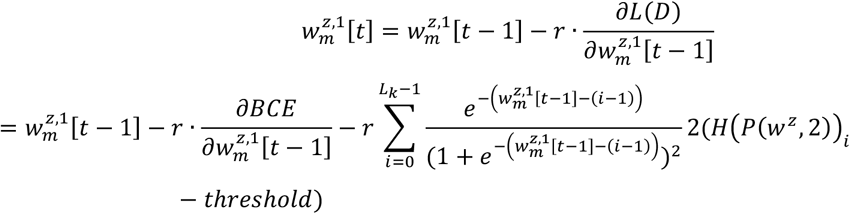

The Shannon entropies of all kernel positions far from the mask boundaries 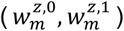 contribute little to the final derivatives. Therefore, if most boundary-flanking kernel positions have a low Shannon entropy (i.e., a high information content; defined by *H*(*P*(*w*^*z*^, 2)) − *threshold* < 0), then the masked Shannon loss (MSL) will help push the boundaries outwards (i.e.,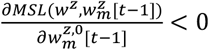 and 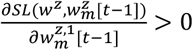 during gradient descent, just as if the current mask is too narrow to span all informative positions. Likewise, if most boundary-flanking kernel positions have a high Shannon entropy (i.e. a low information content; defined by *H*(*P*(*w*^*z*^, 2))_*i*_ − *threshold* > 0), then the MSL will help push the boundaries inwards (i.e.,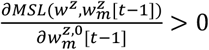 and 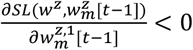), just as if the current mask is too broad to exclude some positions with low information content.

To speed up computation in practice, we approximate the sum by ignoring most small, outside-mask kernel positions and retaining only those terms with 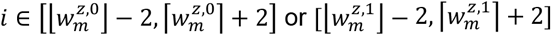.

Finally, when introducing a vConv layer into a model, values of 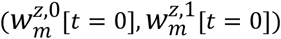, and *threshold* in MSL are of user’s choice. In all subsequent benchmarking on vConv-based networks, unless otherwise specified, we set for each vConv layer the 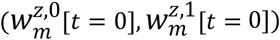 as 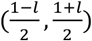 for an unmasked kernel length of *l*, the *λ* as 0.0025, and the *threshold* in MSL as 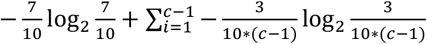 for a kernel with *c* channels. This value of *threshold* is the Shannon entropy for the (discrete) distribution whose highest probability is 0.7, and all others *c*-1’s are equally divided. When processing the nucleic acid sequence, there are totally *c=*4 different probability values, so the probability of the remaining 3 items equally divided is (1-0.7)/3; on the other hand, in Basenji-based Basset networks, the *c* for deeper convolutional layers is not necessarily 4.

### Benchmark vConv-based networks on simulated sequence classification

We simulated for each motif case 6,000 sequences (with 3,000 positive and 3,000 negative) of length 1,000, picked 600 as the test dataset randomly, and split the rest into 4,860 training and 540 validation sequences by setting “validation_split=0.1” in model.fit of Keras Model API (version 2.2.4) [41]; in this way, for each motif case, the same collection of training, validation, and test datasets was used for all hyper-parameter settings tested (see below). The ratio of counts of positive to negative sequences in the test dataset was within [0.8, 1.2] for all 7 motif cases. All sequences were generated independently from each other. Each negative sequence is a random sequence whose bases were independently sampled from the categorical distribution P(A)=P(C)=P(G)=P(T)=0.25. For each positive sequence, we first constructed a random sequence as described above, then overwrote/replaced a random-choice fragment with a sequence fragment generated from one of the motifs associated with the case in question (“signal fragment”, see Table 1 for more details). For the cases of 2, 4, 6 and 8 motifs, new motifs were introduced incrementally (i.e., all motifs in “2 motifs” were also included in “4 motifs”, all motifs in “4 motifs” were also included in “6 motifs”, and all motifs in “6 motifs” were also included in “8 motifs”).

**Table 1.**
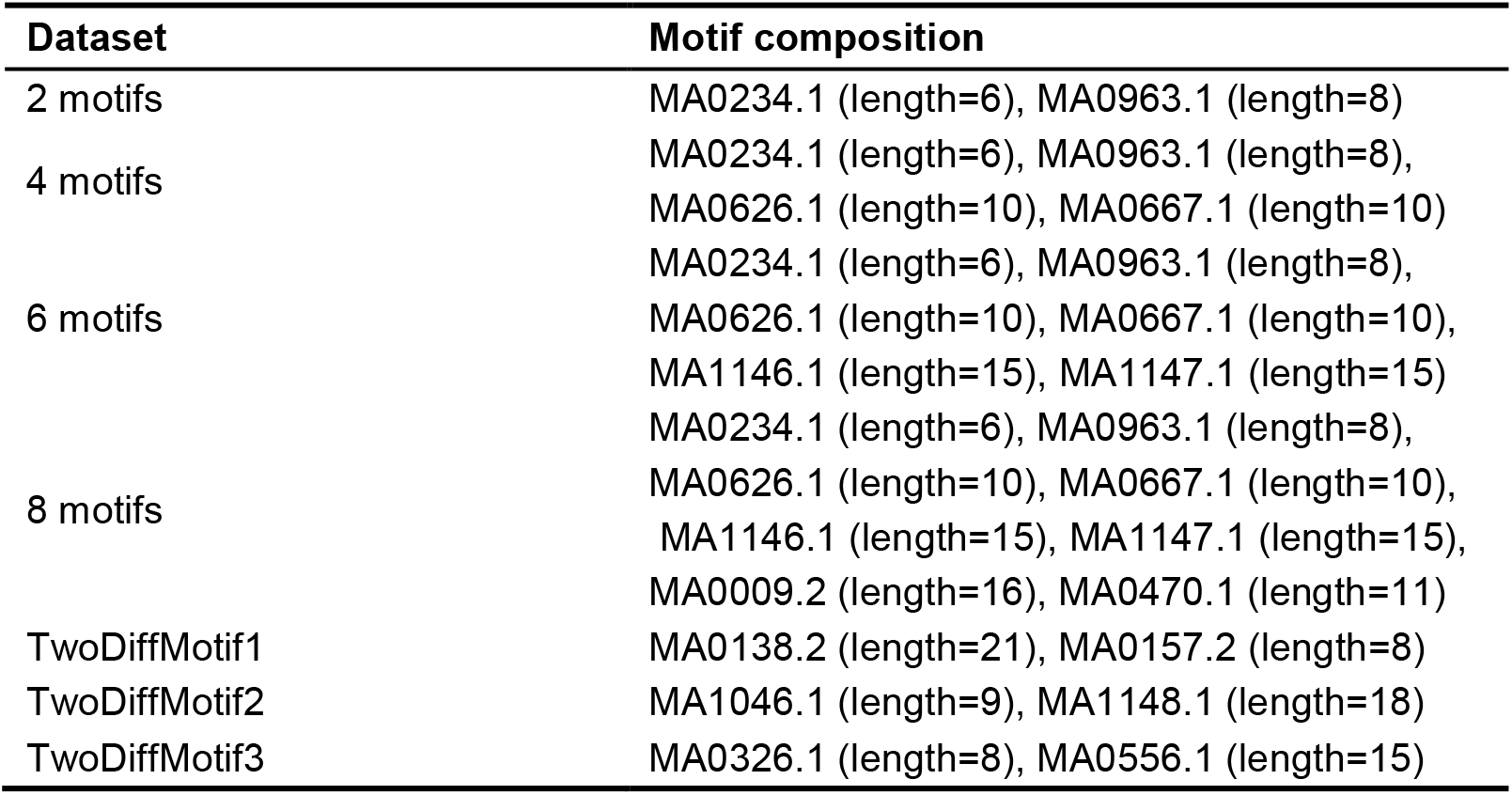
Motifs used to generate each dataset. All motifs were derived from JASPAR [42].

In each iteration of benchmarking a certain case, we (1) picked a hyper-parameter setting from Table 2; (2) used the training and validation datasets to train first a vConv-based network (see Supplementary Fig. 1 for its structure) with this hyper-parameter setting and then a canonical convolution-based network with a model structure identical to that of the vConv-based network, except that the vConv layer was replaced with a canonical convolutional layer with the same kernel number and (unmasked) kernel length; and (3) obtained the AUROC values of these two networks on the test dataset. Iterating over all possible hyper-parameter settings yielded a series of AUROC values for both the vConv-based network and the convolution-based network for the case at hand.

**Table 2.** Hyper-parameter space for the comparison between vConv-based networks and convolution-based networks on simulation datasets.

| Hyper-parameter | Value range space |
| --- | --- |
| Kernel number | {64, 96, 128} |
| Kernel length | {6, 8, 10, 12, 14, 16, 18, 20} |
| Random Seeds | 16 different seeds |
| Momentum in AdaDelta optimizer | {0.9, 0.99, 0.999} |
| Delta in AdaDelta optimizer | {1e-4, 1e-6, 1e-8} |

We used AdaDelta [43] with learning rate 1 as the optimizer. All parameters except for vConv-exclusive ones were initialized by the Glorot uniform initializer [44]; vConv-exclusive parameters were initialized as specified in the subsection “Design and implementation of vConv”. Both convolution-based and vConv-based networks were trained with early stopping on validation loss with patience set to 50. All vConv-based networks with *n* kernels except for vConv-based Basset networks were first trained without updating 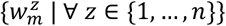 for 10 epochs and then trained with updating 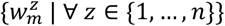 for vConv-based Basset networks, we updated 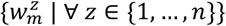 since the first epoch.

### Benchmark vConv-based networks on real-world sequence classification

To further demonstrate the performance of vConv-based networks in the real world, we examined whether replacing convolutional layers with vConv helps to improve the performance for motif identification in real-world ChIP-Seq data, against three published convolution-based networks: DeepBind [26], Zeng et al.’s convolution-based networks [45], and Basset [46].

For the first comparison, we downloaded DeepBind’s original training (with positively sequences only) and test datasets from https://github.com/jisraeli/DeepBind/tree/master/data/encode, and generated negative training dataset by shuffling positive sequences with matching dinucleotide composition as specified by DeepBind [26]. We then trained vConv-based networks (see below for the model structure and training details) and compared their AUROC on test datasets with those of three sets of DeepBind performances from Supplementary Table 5 of DeepBind’s original paper [26]: (1) the DeepBind* networks, which were trained by using all the training dataset; (2) the DeepBind networks, whose positive training sequences consist of those from the top 500 odd-numbered peaks only; and (3) the DeepBindbest networks, the one of DeepBind and DeepBind* with larger AUROC for each test dataset. During comparison, we found that there are two datasets with different AUROC values yet sharing the same model name in their original dataset (REST_HepG2_NRSF_HudsonAlpha and REST_SK-N-SH_NRSF_HudsonAlpha), which makes us unable to determine their true AUROC values, so we excluded them from the comparison.

For the second comparison between vConv-based networks and Zeng et al.’s [45] convolution-based networks, we downloaded 690 ENCODE ChIP-Seq-based training and test datasets representing the DNA binding profile of various transcription factors and other DNA-binding proteins from http://cnn.csail.mit.edu/motif_discovery/. We then trained vConv-based networks (see below for the model structure and training details) and compared its AUROC value on test datasets with the AUROC values of Zeng et al.’s networks from http://cnn.csail.mit.edu/motif_discovery_pred/.

The first two comparisons used the same model structure (Supplementary Fig. 1) and hyper-parameter space (Supplementary Table 1). For direct comparison between vConv-based networks and convolution-based networks for DeepBind and Zeng et al.’s cases [45], the vConv-based network was implemented by replacing the convolutional layer of each network considered for comparison with a vConv layer (with exactly the same hyper-parameters for the number of kernels and the initial (unmasked) kernel length). The hyper-parameter space is listed in the Supplementary Table 1. The remaining details of parameter initialization followed those of the corresponding convolution-based networks [26, 45], and all these networks used the training strategy described in the simulation case.

Finally, for the third comparison with the Basset network [46], we used the Basenji reimplementation of Basset [28] (https://github.com/calico/basenji/tree/master/manuscripts/basset) made and recommended by the original author of Basset (Kelley D., personal communications); this reimplementation is a deep CNN with nine convolutional layers (see Supplementary Fig. 2 for the full details). We (1) generated the datasets by running “make_data.sh“ under “manuscripts/basset/” (of this repository), (2) re-trained the original Basset networks (which makes prediction for 164 cell types by a single model) by running “python basenji_train.py -k -o train_basset params_basset.json data_basset” under “manuscripts/basset/” (of this repository) to obtain their AUROC values, (3) replaced either the first (‘Single vConv-based’; left figure in Supplementary Fig. 3) or all nine (‘Completed vConv-based’; right figure in Supplementary Fig. 3) convolutional layers in the Basset network with vConv layer(s), re-initialized all weights, and retrained it to obtain the AUROC value of the vConv-based Basset network, and finally (4) compared the AUROC values between the two types of networks. The details of parameter initialization (except for the vConv-exclusive parameters) and stochastic gradient descent-based optimization (with learning rate 0.005) were exactly the same between the vConv-based and the original Basset networks (Supplementary Fig. 2 and 3).

### Benchmark vConv-based networks on real-world motif discovery

Finally, we compared vConv-based networks with the canonical methods on the motif discovery problem. We followed the protocol proposed by Zhang et al. (Zhang, et al., 2015) to assess the accuracy of the extracted representative motifs based on the ENCODE CTCF ChIP-Seq datasets [47]. In brief, for each motif discovery tool and each ChIP-Seq dataset, we located the motif-containing sequence fragments for candidate motifs discovered by this tool, checked whether they overlapped with the ChIP-Seq peaks with CisFinder (retrieved from https://lgsun.grc.nia.nih.gov/CisFinder/download.html at Feb 22, 2020), and reported the largest per-motif ratio of overlapping fragments to all fragments across all candidate motifs as the final accuracy of this tool on this dataset (see Supplementary Fig. 4 for more details). For the sake of demonstrating the advantage of vConv-based networks, here the model structure of the vConv-based network was chosen to be the one used in simulation (Supplementary Fig. 1).

## RESULTS

### vConv-based networks identify motifs more effectively than convolution-based networks

We first present direct comparisons between vConv-based and convolution-based networks based on multiple simulated datasets (see the Methods for more details). vConv-based networks performed statistically significantly better across most hyperparameter settings of all seven cases (Fig. 2; with all Wilcoxon rank sum test’s p-values < 1e-18 for the null hypothesis that the AUROC of the vConv-based network is equal to or smaller than that of the convolution-based network). This improvement became larger when more motifs are introduced (2 motifs to 8 motifs in Supplementary Fig. 5) and with increased heterogeneity of the motif length (2 motifs v.s. TwoDiffMotif1/2/3 in Supplementary Fig. 5). In addition, vConv-based networks showed a smaller mean standard error of the AUROC than convolution-based networks across different hyper-parameter settings (Levene’s test, p-value < 0.001 for all datasets; see Supplementary Table 2), suggesting that vConv-based networks also have a better robustness to hyper-parameters than convolution-based networks.

**Fig. 2.**
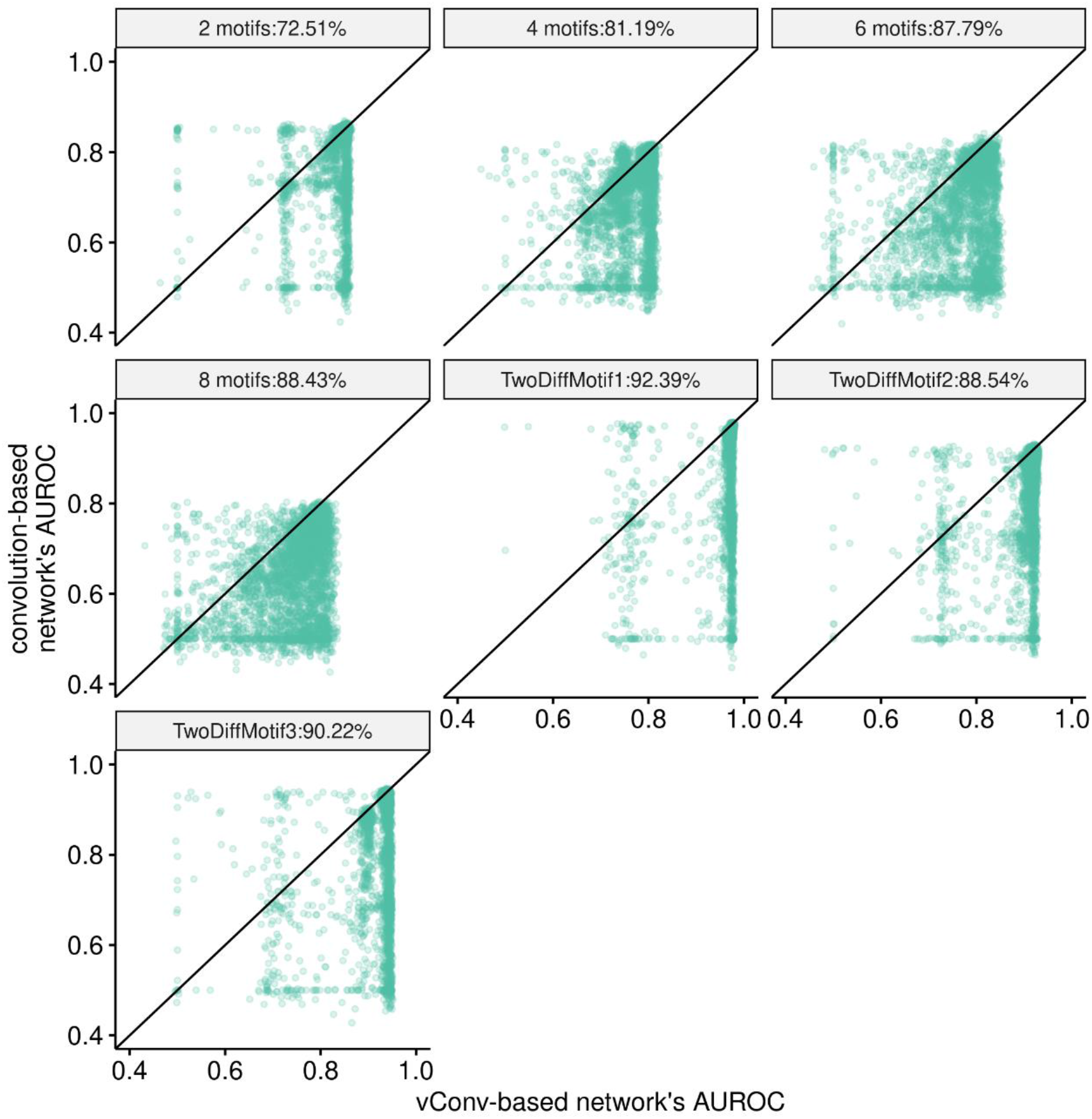
Pairwise comparison of AUROC between vConv-based and convolution-based networks with exactly the same hyper-parameter. As expected, vConv-based networks performed better than convolution-based networks in most cases. The percentage in the title on the figure indicates the percentage of those cases where the vConv-based network was better than the convolution-based network.

We further visualized the first layer of trained vConv-based and convolution-based networks on the most complex dataset (8 motifs), and found that vConv-based networks accurately recovered the underlying real motif from sequences and converge to the real length well (Supplementary Fig. 6). Of note, we found that the vConv-based networks successfully learned a motif (MA0234.1) which was missed by canonical convolution-based networks (Supplementary Fig. 6, second row).

These findings further compelled us to suspect that vConv-based networks will perform better than convolution-based networks on real-world cases possibly with combinatorial regulation.

To test this hypothesis, we examined whether replacing convolutional layer(s) with vConv would improve performance of the following convolution-based networks: (1) DeepBind [26], a series of DNA-protein-binding classifiers trained for each of 504 ENCODE ChIP-Seq datasets; (2) Zeng et al.’s convolution-based networks [45], a series of structurally optimized variants of DeepBind models for each of 690 ChIP-Seq datasets; and (3) Basset [46], a complex DNase footprint predictor with nine convolutional layers (see Methods for more details). As expected, vConv-based networks showed a statistically significantly improved performance compared to their convolution-based network counterparts for all these three cases (Fig. 3 and Supplementary Fig. 7), with a few exceptions some of which can be attributed to small sample size of the input dataset (Supplementary Fig. 8A-B) and disappeared upon additional grid-search on kernel length (Supplementary Fig. 8C). Of note, stacking vConv layers for a complex convolution-based network like Basset can statistically significantly improve the performance with respect to both the original Basset network and the single vConv-based network (Fig. 3C), well demonstrating its combined power with the popular hierarchical representation learning strategy in CNN application.

**Fig. 3.**
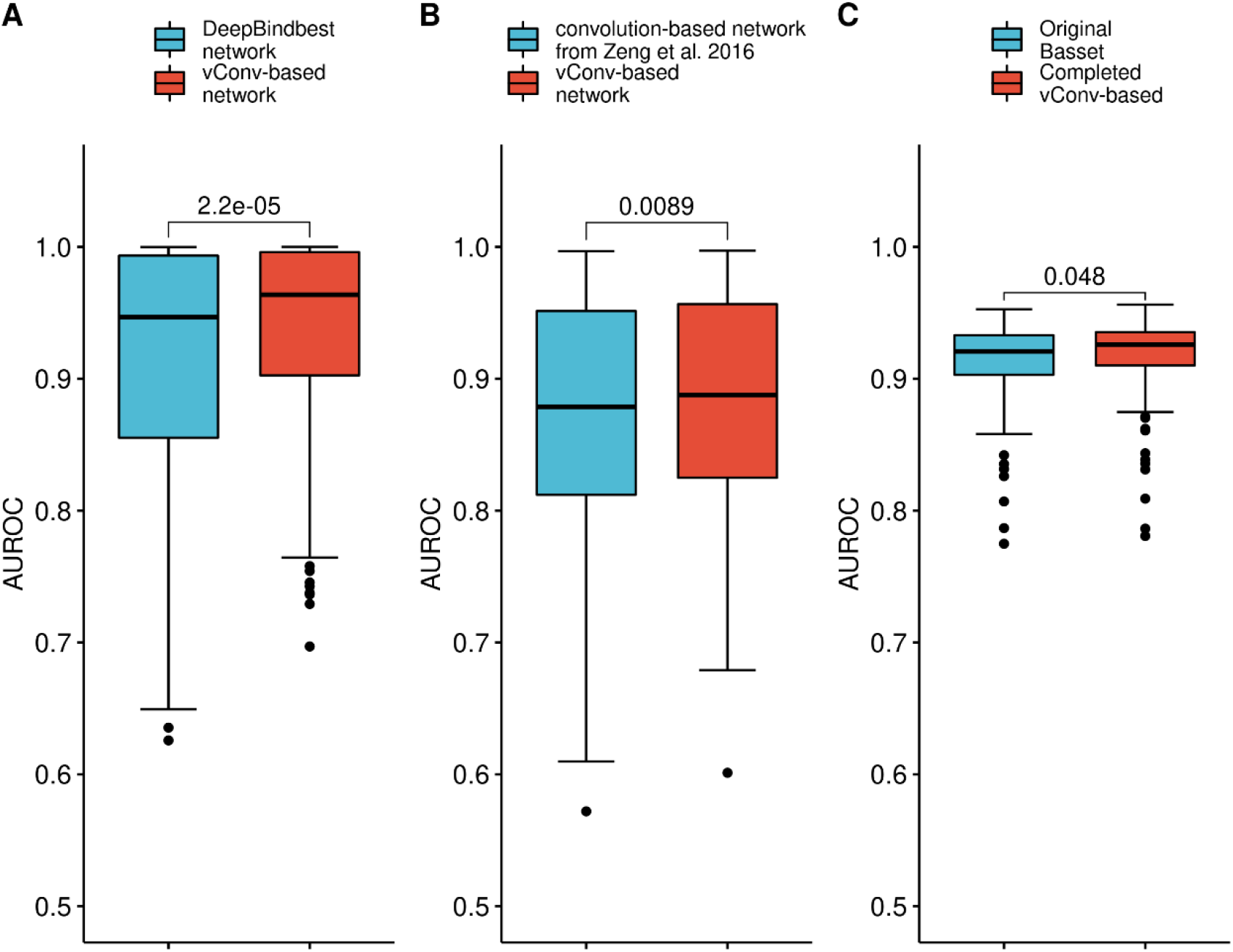
vConv-based networks outperformed convolution-based networks on real-world cases. (A), comparison of AUROC between vConv-based networks and DeepBindbest networks for DeepBind (see also Supplementary Fig. 7 for comparison with other DeepBind networks). (B), comparison of AUROC between vConv-based networks and networks from Zeng et al. . (C), comparison of AUROC between vConv-based networks with ‘Completed vConv-based’, the vConv-based network with all convolutional layers of Basset network replaced by vConv (right figure in Supplementary Fig. 3; see Methods for more details) and networks from Basset. All p-values shown are from Wilcoxon rank sum test, single-tailed, with the null hypothesis that the AUROC of the vConv-based network is equal to or smaller than that of the convolution-based network. See Methods for more details of each specific network.

### vConv-based networks discover motifs from real-world sequences more accurately and faster than canonical tools

We further evaluated vConv-based networks’ performance for *ab initio* motif discovery. In brief, for a particular trained vConv-based network, we first selected kernels with corresponding dense layer weights higher than a predetermined baseline (defined as mean (all dense layer weights) - standard deviation (all dense layer weights)) and then extracted and aligned these kernels’ corresponding segments to compute the representative PWM (Fig. 4A). We then compared the accuracy of recovering ChIP-Seq peaks by the vConv-based motif discovery and other motif discovery tools across all these ChIP-Seq datasets [47] (see the Methods for details).

**Fig. 4.**
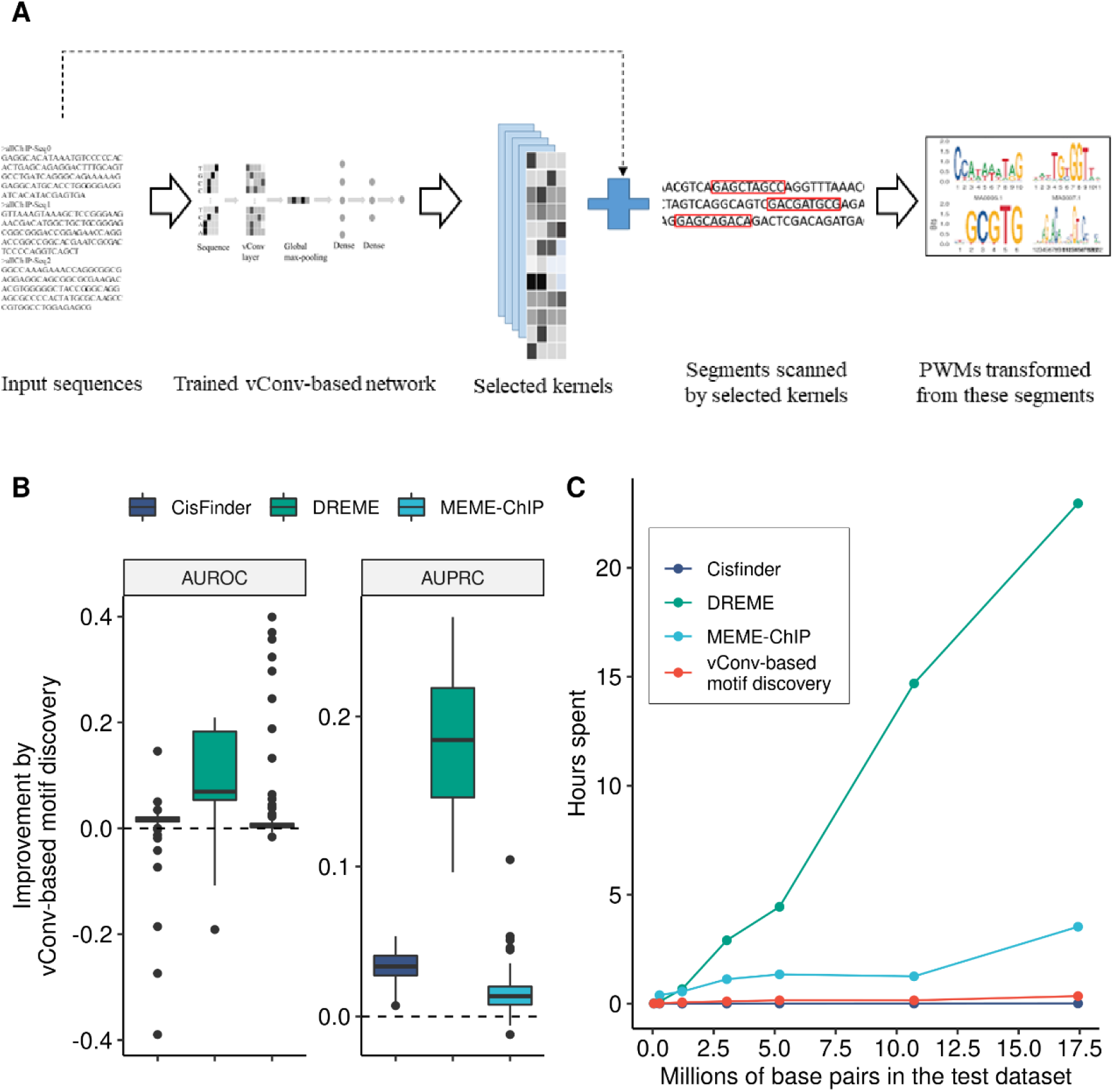
vConv-based networks discover motifs more accurately and faster. (A) shows the process of calling a representative motif from a trained vConv-based network. (B) shows the difference in accuracy, defined as the accuracy of vConv-based motif discovery minus the accuracy of the motif discovery algorithm shown on the x-axis on the same dataset. MEME [24] failed to complete the discovery within a reasonable time (∼50% datasets remained unfinished even after running for 1.5 weeks with 2,000 cores, amounting to 504,000 CPU hours) for these datasets, and its results were thus not listed here. See also Supplementary Fig. 9 for other metrics. (C) shows the time cost of each motif discovery algorithm as a function of millions of base pairs in the dataset tested.

Out of all 100 CTCF datasets, the vConv-based network outperformed DREME [15] on 95 datasets, CisFinder [23] on 91 datasets and MEME-ChIP [25] on 87 datasets with respect to AUROC (Fig. 4B; see also Supplementary Fig. 9 for other metrics). In addition, the vConv-based network ran much faster than DREME and MEME-ChIP on large datasets (more than 17 million base pairs finished in less than 30 minutes compared to hours by DREME and MEME-ChIP’s; see Fig. 4C).

## DISCUSSION

Inspired by the theoretical model (Supplementary Notes 1) for analyzing the influence of the kernel size in the convolutional layer, we designed and implemented a novel convolutional layer, vConv, which adaptively tunes the kernel lengths at run time. Evaluations based on both simulations and real-world datasets showed that such design enables vConv-based networks to outperform canonical convolution-based network (see Fig. 3 and 4, as well as Supplementary Fig. 5-8 for more details), especially on datasets containing multiple combinatorially regulatory *cis*-elements of various lengths [38, 48]. Meanwhile, vConv could be readily integrated into multi-layer neural networks, as an “in-place replacement” of canonical convolutional layer with potential applications in more scenarios such as *cis*-regulatory motif prediction [49], predicting non-coding functions *de novo* [50], predicting the transcription factors binding intensities [51], and quantitative detection of histone modifications [52]. To facilitate its application in various fields, we have implemented vConv as a new type of convolutional layer in Keras (https://github.com/gao-lab/vConv).

In theory, a regular fixed-length convolution kernel with size larger than that of the *bona fide* motif, may be able to converge to a solution where the kernel captures the motif faithfully with zero padding everywhere else, just as what the vConv kernel does. Nevertheless, preliminary empirical evaluations suggest that such “perfect” solution could be very rare in reality with a general training strategy: none of the convolution-based network kernels similar to the ground truth motifs in the 8 motifs case (as determined by Tomtom; see Supplementary Fig. 6) contains more than 1 position whose sum of absolute elements are one order of magnitude more closer to 0 than other positions.

vConv is scalable considering the fact that its current design only adds a *O(ncl)* per *n*-kernel-convolutional layer (with kernel length *l* and number of channels *c*) replaced to the overall time complexity compared with that of the canonical fixed-kernel convolutional layer, and that preliminary empirical evaluations suggest no evidence of convergence slowdown (as benchmarked in Supplementary Fig. 10 for the simulation case). While we notice that the current variable-length kernel strategy implemented by vConv could, in theory, be approximated by combing multiple fixed-length-kernel networks, we’d argue that such approximation would lead to serious inflation for the number of trainable parameters, and higher time cost for training and optimization. For example, to approximate a simple network with one vConv layer of *n* unmasked kernels with average length *l* (which needs *n*\*(4\**l*+2)+*n* = *n*\*(4\**l*+3) parameters), the ensemble of *m* convolution-based networks would take *m*\*(*n*\*(4\**l*)+*n*) = *m*\*(*n*\*(4\**l*+1)) parameters, far larger than *n*\*(4\**l*+3) when *m* is of tens or hundreds - a popular choice in ensemble learning.

vConv-based networks show robustness with various hyper-parameters and initialization setups (Supplementary Table 2). We believe that the major vConv-exclusive components, Shannon loss (MSL) and boundary parameters, contribute to the increased robustness. Intuitively, such components could regularize the parameters to learn, making the overall objective function simpler (i.e., having fewer suboptima) and thus more robust. Consistently, we found out that the vConv-based network -- when equipped with MSL -- had a smaller variance of AUROC that is statistically significantly different from that of the vConv-based network without MSL in 3 out of all 7 simulation cases (Levene’s test, p<0.001; Supplementary Table 3), suggesting that MSL does contribute to this increased robustness in certain conditions. Of interest, equipping MSL also improved the AUROC of the vConv-based network (Supplementary Fig. 11).

## FURTHER WORK

We note that the current theoretical analysis (see details in Supplementary Notes 1) reported above relies on prior knowledge of the real motif ℳ, which may not be feasible to obtain for real-world datasets. A possible workaround is to estimate the empirical null distribution of P_*real*_ over a set of randomly initialized PWMs and kernels and then derive the expectation of P_*real*_ over the given dataset. Such theoretical distribution, once established, could serve as a general background distribution for further development of motif scanners.

Moreover, although current motifs in convolution-based networks are automatically represented by PWMs, this might be an oversimplified representation of the genuine motifs, which may allow insertions and deletions within motifs (e.g., the HMM motifs from Pfam [53] and Rfam [54]). While recurrent neural networks have been expected to be able to learn such motifs [55-58], the interpretation of such models is still challenging. A promising alternative would be to use the CNN framework to learn complex motifs directly, as demonstrated by a recently developed CNN model HOCNNLB [59] which can learn first-order Markov models by re-encoding input sequences. Combining the variable-length nature of vConv and such complex-motif-capturing models might be much more useful for mining biological motifs.

## DATA AVAILABILITY

A Keras-based implementation is released at GitHub (https://github.com/gao-lab/vConv) for academic usage, with all scripts/data for reproducing results available at https://github.com/gao-lab/vConv-Figures_and_Tables.

## ACKNOWLEDGEMENT

The authors would like to thank Drs. Cheng Li, Letian Tao, Minghua Deng, Zemin Zhang, Jian Lu and Liping Wei at Peking University for their helpful comments and suggestions during the study. The analysis was supported by the High-performance Computing Platform of Peking University, and we thank Dr. Chun Fan and Yin-Ping Ma for their assistance during the analysis.

## KEY POINTS

- We proposed a novel convolutional layer, vConv, which learns kernel structure from data adaptively.
- vConv could be readily integrated into multi-layer neural networks, as an “in-place replacement” of canonical convolutional layer.
- vConv’s high robustness and low time footprint further enable its usage for real-world massive omics data.

## FUNDING

This work was supported by funds from the National Key Research and Development Program [grant number 2016YFC0901603]; the China 863 Program [grant number 2015AA020108]; the State Key Laboratory of Protein and Plant Gene Research; the Beijing Advanced Innovation Center for Genomics (ICG) at Peking University; and in part by the National Program for the Support of Top-notch Young Professionals [to G.G.].

## Conflict of interest statement

None declared.

## Notes

https://github.com/gao-lab/vConv

### Competing Interest Statement

The authors have declared no competing interest.

