## Supplementary material for "Identifying complex motifs in massive omics data with a variable-convolutional layer in deep neural network": Supplementary_figures_1-6.pdf

**SUPPLEMENTARY FIGURES -- FIRST PART**

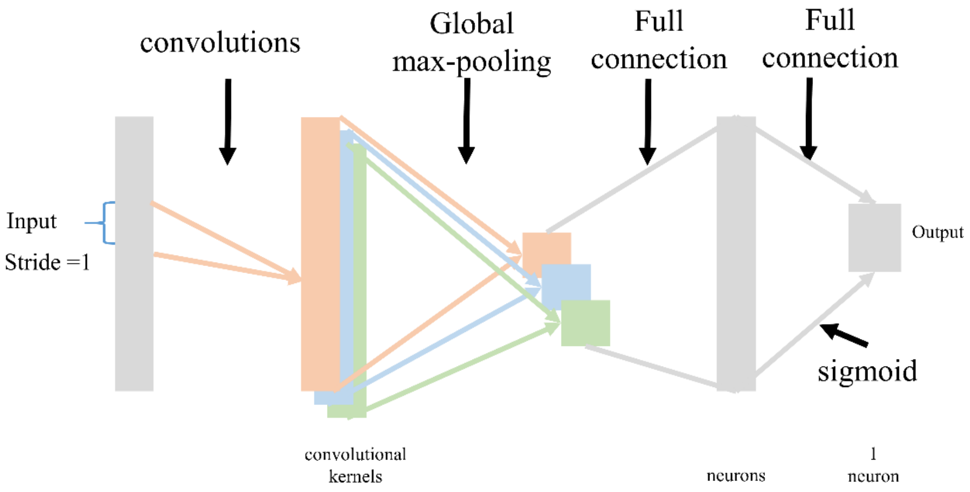


**Supplementary Fig. 1.** The model structure of vConv-based networks used in benchmarking with simulation, DeepBind, and Zeng et al.'s convolution-based networks as well as the one used by the vConv-based network in comparison of motif discovery.

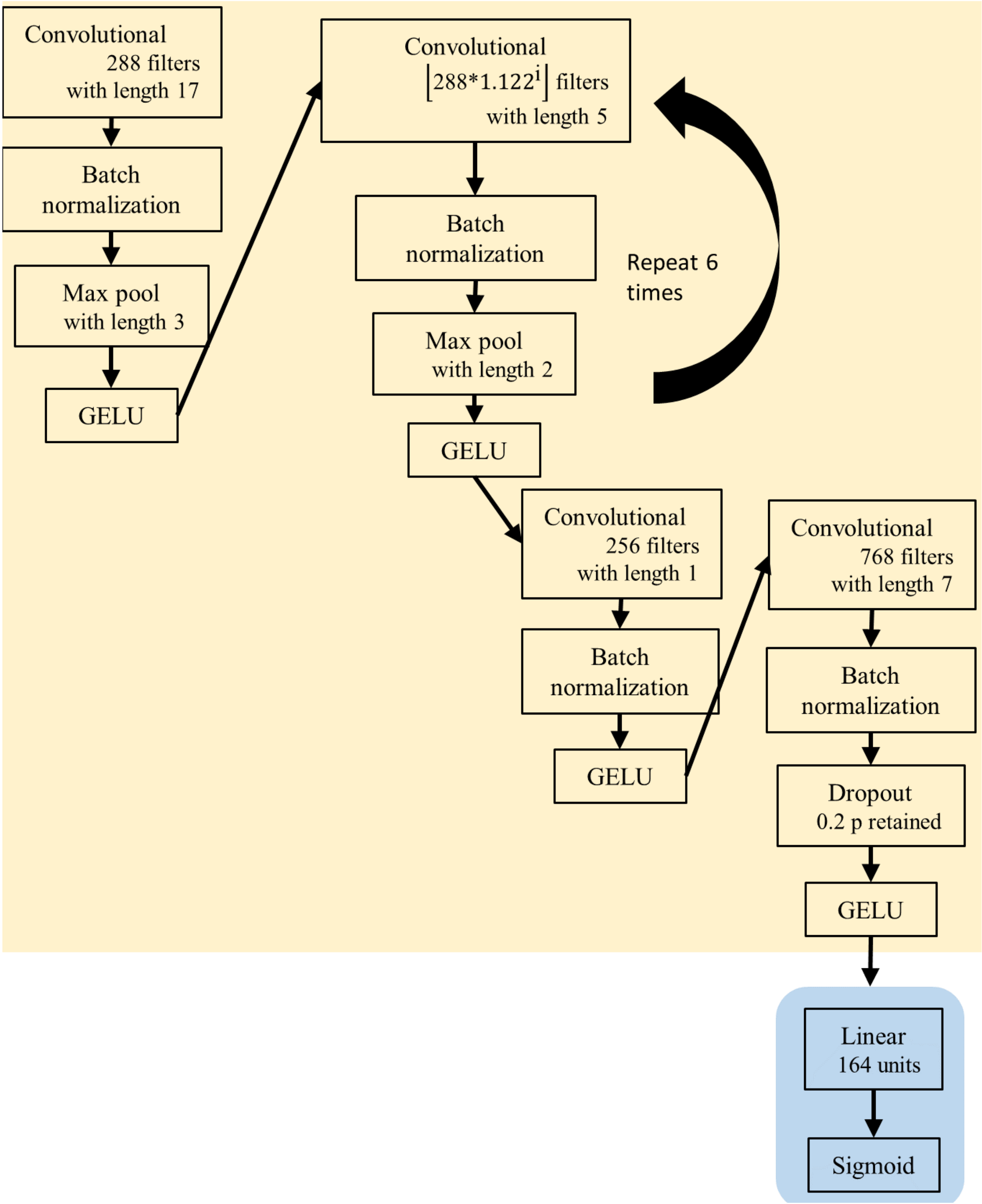


**Supplementary Fig. 2.** The detailed view of the Basenji-based Basset network that we studied throughout the manuscript**.** The parameter i in figure is the increasing order of this layer among all 6 repeats, starting from 1.

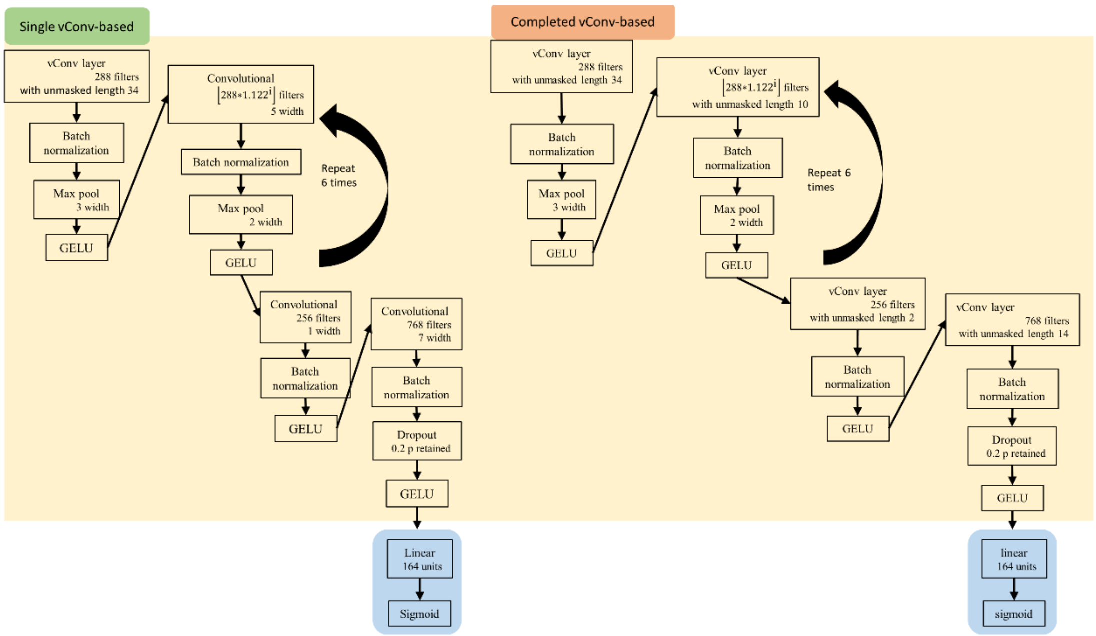


**Supplementary Fig. 3.** The detailed view of vConv-based Basset networks that we studied throughout the manuscript. The network in the left ('Single vConv-based') figure replaced only the first convolutional layer with a vConv layer, while the network in the right ('Completed vConv-based') replaced all the convolutional layers. The parameter i in figure is the increasing order of this layer among all 6 repeats, starting from 1.

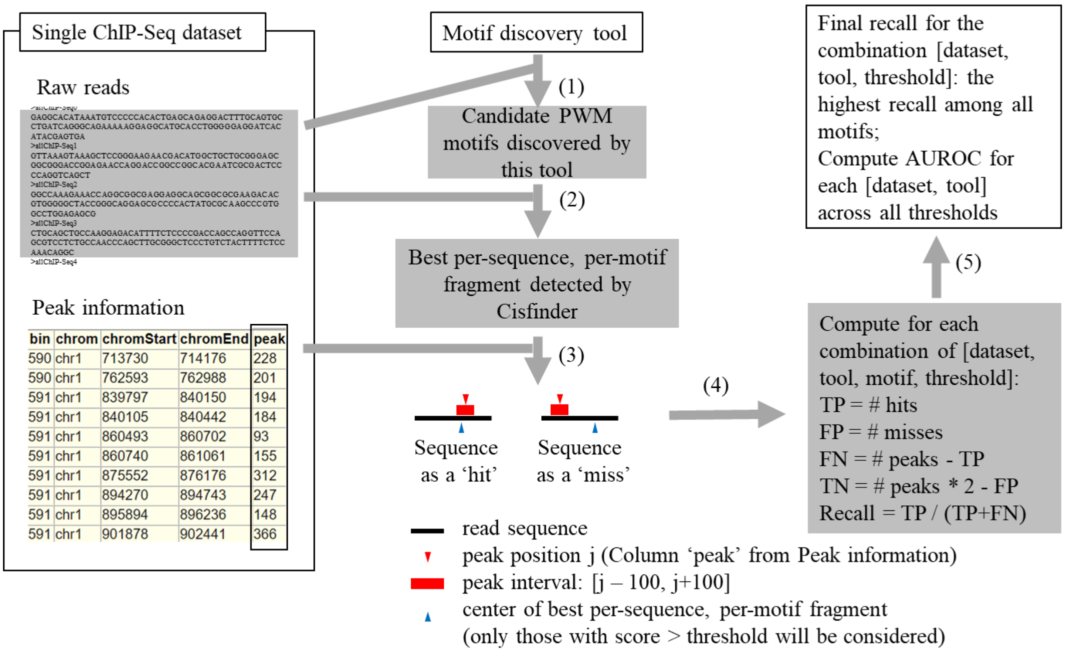


**Supplementary Fig. 4.** The pipeline for computing the recall of recovery of ChIP-Seq peaks using a specific motif discovery tool on a specific ChIP-Seq dataset. In brief, (1) the motif discovery tool was used on raw reads of the ChIP-Seq dataset to produce candidate PWM motifs; (2) patternScan from CisFinder [1] was used to identify, for each raw read sequence and each PWM motif, the sequence fragment that is most likely to be generated by the PWM (we used a single motif instead of all motifs output by the model as the input to patternScan because patternScan will merge similar sites for all motifs, causing some motifs to lose their corresponding sites); (3) it was determined whether each sequence fragment overlaps with a peak interval (defined as [j-100,j+100], where j is the position of the real peak), and a sequence was counted as a hit if its fragment aligned with the genome and overlapped with the peak, or as a miss otherwise; (4) the TP, FP, TN, FN, recall and other metrics for each motif on this dataset was calculated accordingly, following those in MOST+ [2]; and (5) the motif with the highest recall was chosen as the final PWM, and its recall was taken as the final recall.

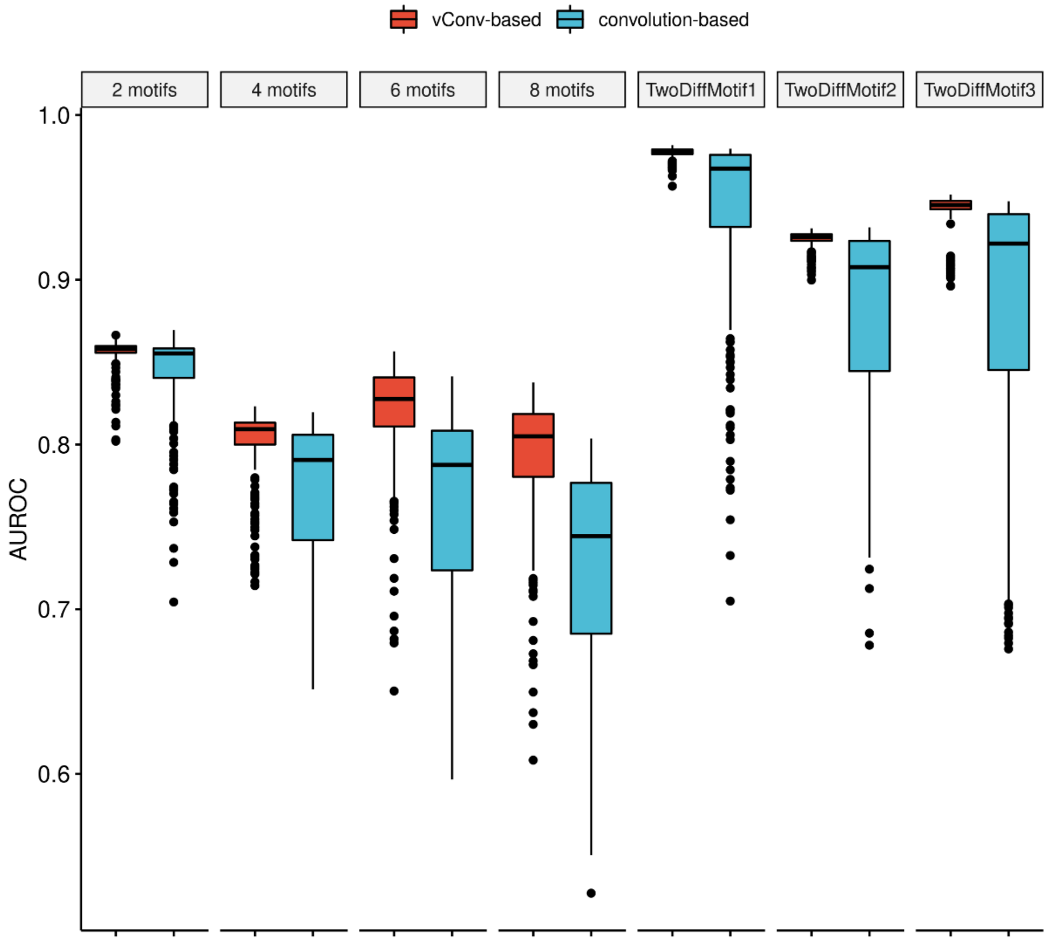


**Supplementary Fig. 5.** vConv-based networks outperformed canonical convolution-based networks in overall AUROC comparison for motifs of different lengths on simulation datasets. For each hyper-parameter settings (excluding the random seed), all AUROC values from all random seeds were considered.

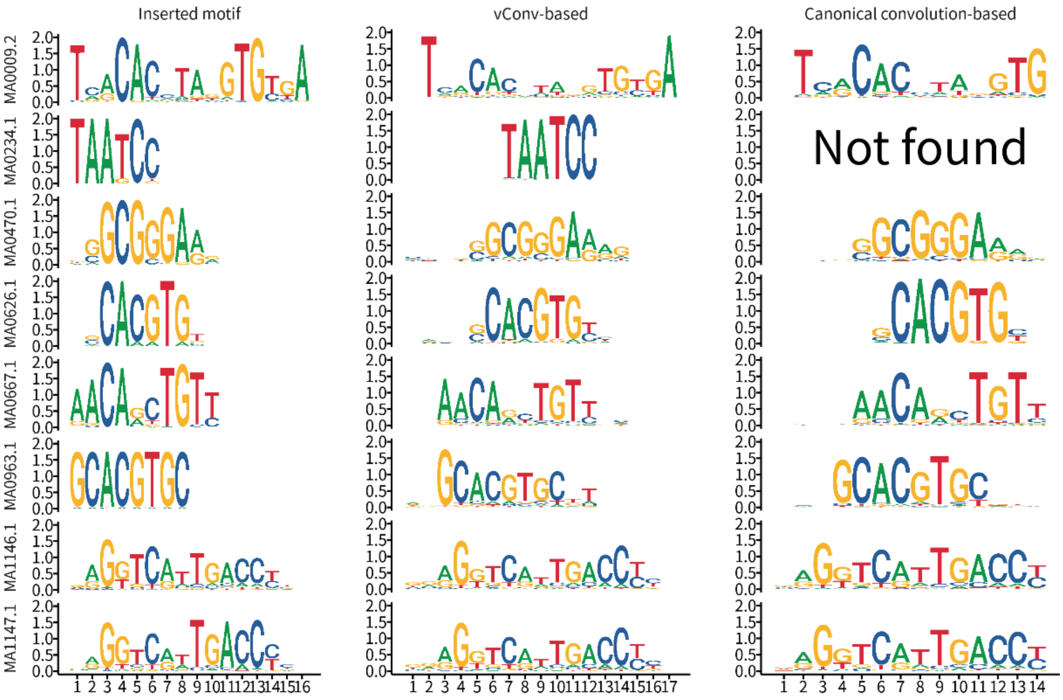


**Supplementary Fig. 6.** The vConv-based networks accurately recovered the underlying real motif from sequences and converge to the real length well in the 8 motifs case. The x-axis denotes base positions along the motif, while the y-axis is the information content for each base position. We used Tomtom v5.3.0[3] to compare whether the PWM of each learnt vConv or convolutional kernel is similar to the ground truth motif inserted into the simulation dataset (details are in canonical_convolutional_tomtom.html and vConv_tomtom.html), and plotted sequence logos for the most similar one (i.e., with the smallest q-value that is less than 0.1). The canonical convolution-based network did not detect any kernel whose PWM has a q-value < 0.1 for the motif MA0234.1.
