## Supplementary material for "Identifying complex motifs in massive omics data with a variable-convolutional layer in deep neural network": Supplementary_figures_7-12.pdf

### SUPPLEMENTARY FIGURES -- SECOND PART

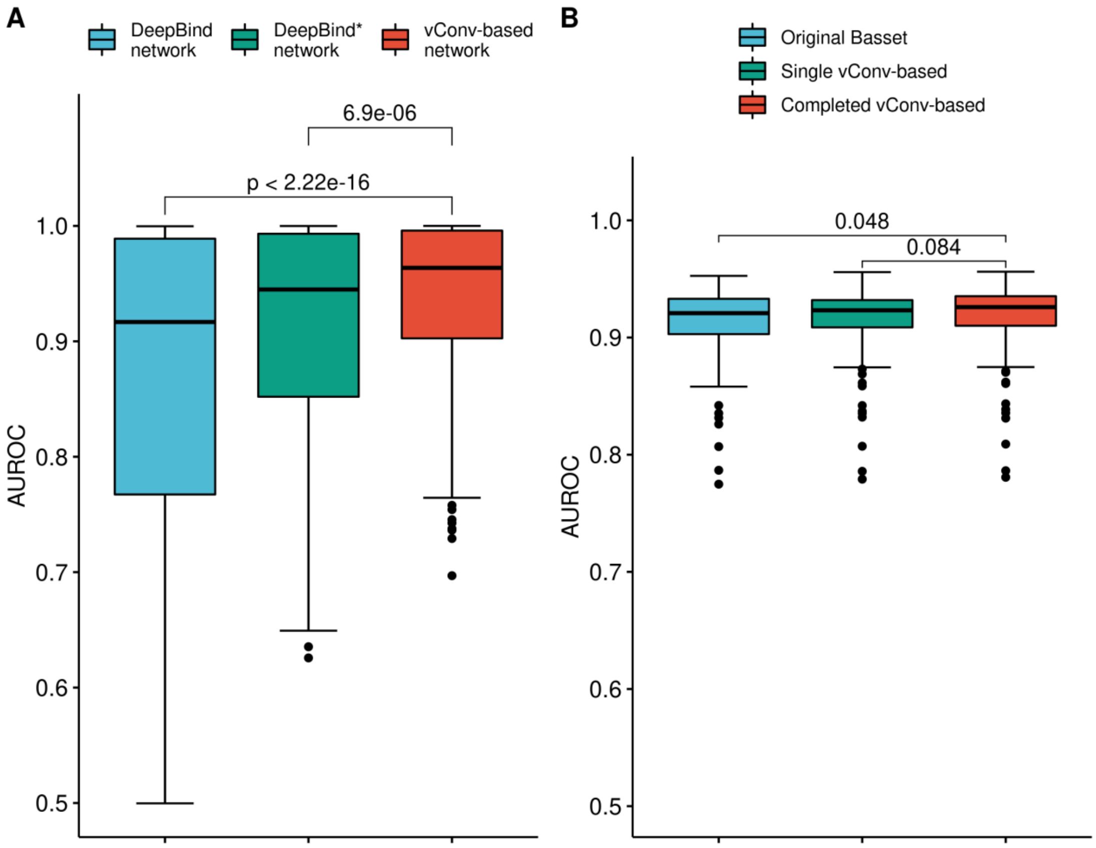


**Supplementary Fig. 7.** Additional comparisons between vConv-based and convolution-based networks. (A), comparison between vConv-based networks, DeepBind, and DeepBind* networks from DeepBind. (B), comparison between the Original Basset, the Single vConv-based network (where only the first convolutional layer of Basset network was replaced with the vConv layer), and the Completed vConv-based network (where all convolutional layers of Basset network were with vConv layers). All p-values shown are from Wilcoxon rank sum test, single-tailed, with the null hypothesis that the AUROC of the vConv-based network is equal to or smaller than that of the convolution-based network.

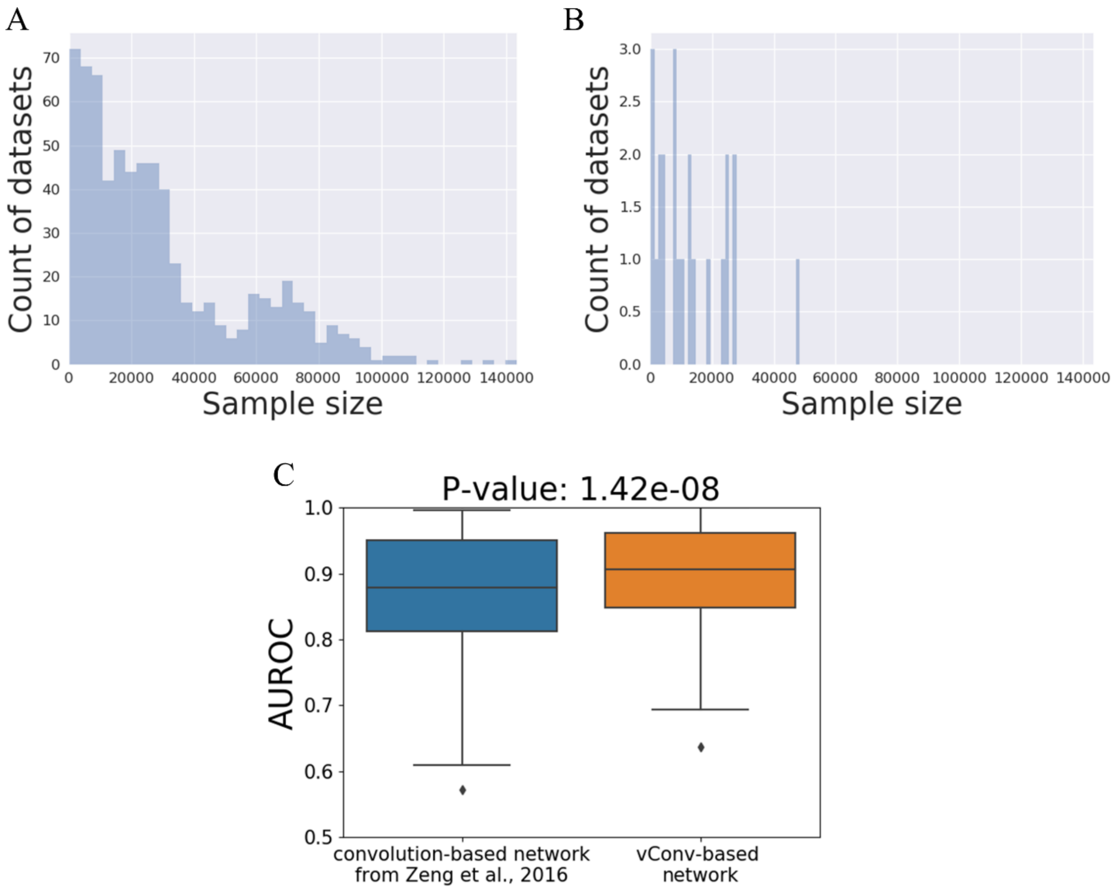


**Supplementary Fig. 8.** Examination of the 23 datasets on which vConv-based networks performed more poorly than Zeng et al.'s convolution-based networks. The sample size distributions were shown for all ChIP-Seq datasets (A) and for the 23 datasets on which the vConv-based network performed more poorly than the convolution-based network (B). A sample size drawn from the distribution in (B) has a probability of 0.70 of being smaller than a sample size drawn from the distribution in (A). When we chose to compare the optimal solution of vConv-based network under the same hyper-parameter space as above compared with DeepBind (see Supplementary Table 1), the vConv-based network performed statistically significantly better than the Zeng et al.'s convolution-based network on these datasets (C). The p-value shown is from Wilcoxon rank sum test, single-tailed, with the null hypothesis that the AUROC of the vConv-based network is equal to or smaller than that of the convolution-based network.

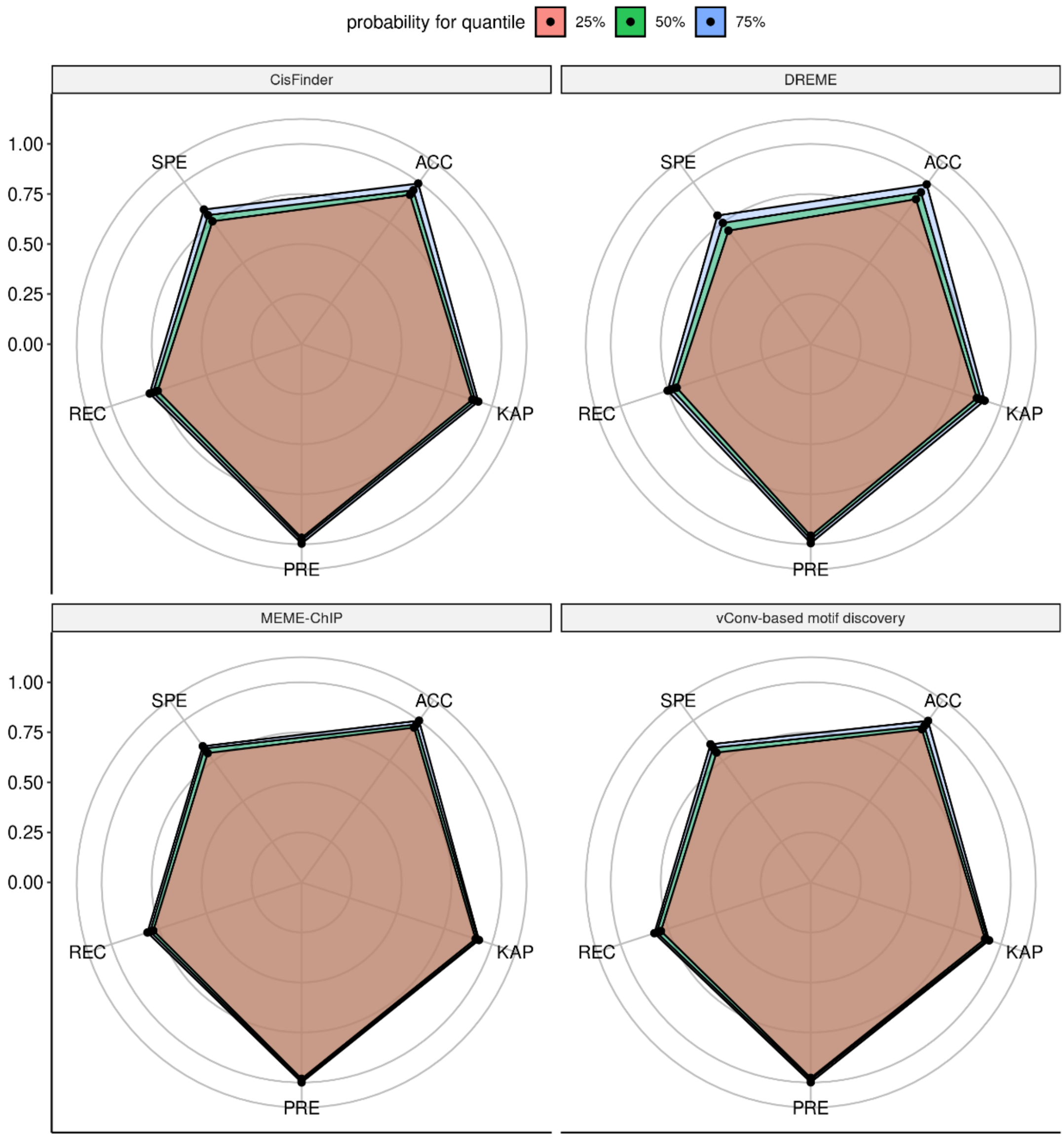


**Supplementary Fig. 9.** Radar plot of five metrics for different tools. Only the 25%, 50%, and 75% quantiles of these metrics across all thresholds are plotted. ACC, accuracy; KAP, kappa; PRE, precision; REC, recall; SPE, specificity.

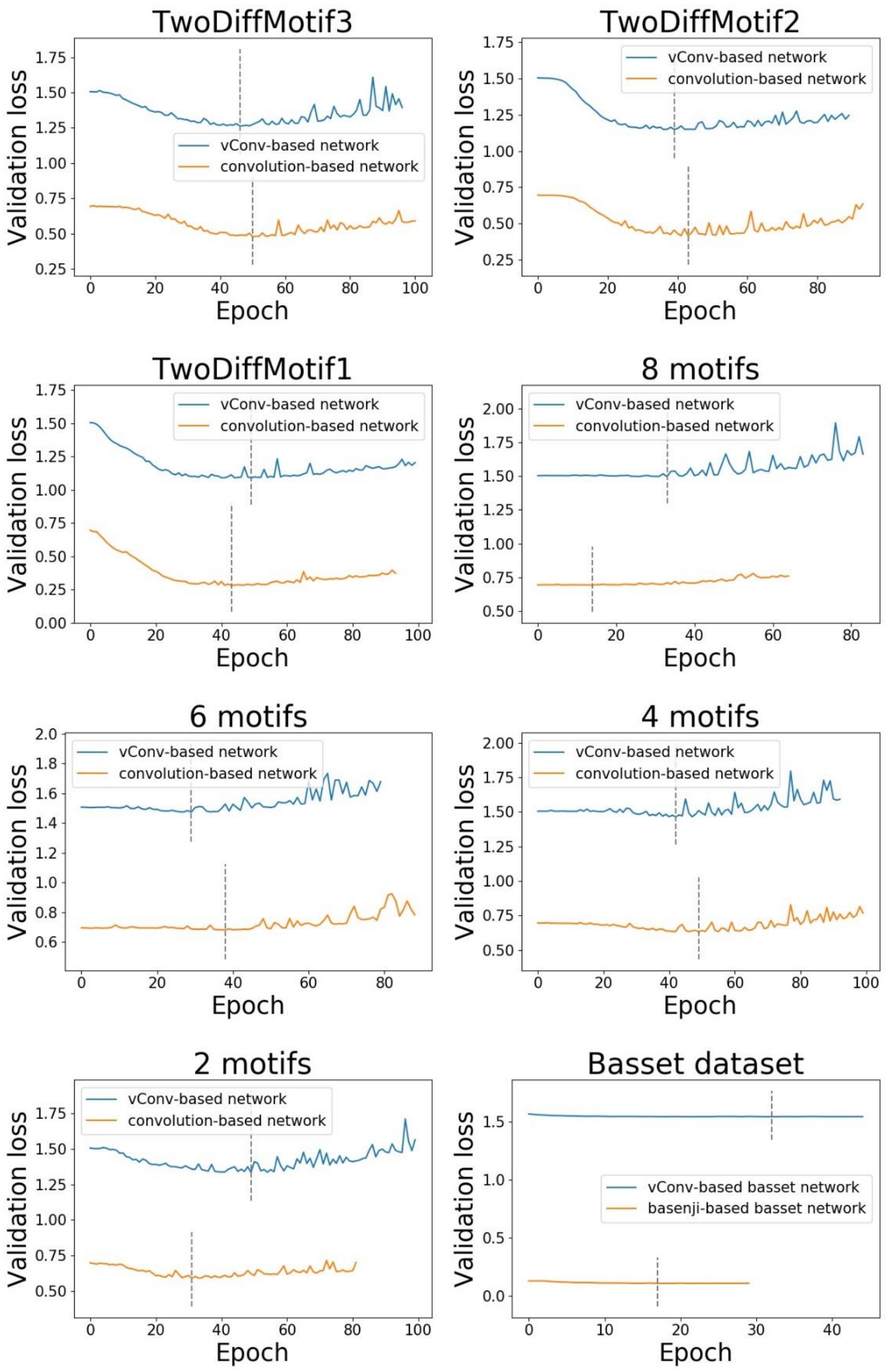


**Supplementary Fig. 10.** Similar speeds of convergence between vConv-based and convolution-based networks on the simulation datasets and Basset datasets. In each subfigure, the x-axis represents the number of epochs, and the y-axis represents the loss on the test subset of the dataset corresponding to that subfigure. The loss function convergence curve we drew includes the epoch after the optimal solution (50 epochs on the simulation data set, and 12 epochs for the basset). The dashed line in the figure is the position corresponding to the optimal solution we used. Because the test and validation subsets were generated from the same distribution, the change in the loss with respect to the number of epochs on the test subset is a good indicator of this change on the validation subset (and thus a good indicator of the speed of convergence). One can see that vConv-based and convolution-based networks converged at a similar speed. Note that the (almost constant) difference in loss is primarily due to the introduction of the Shannon loss by vConv (see the subsection titled 'Design and implementation of vConv' in the Methods section) rather than the poorer performance of the vConv-based network itself (see also Fig. 2).

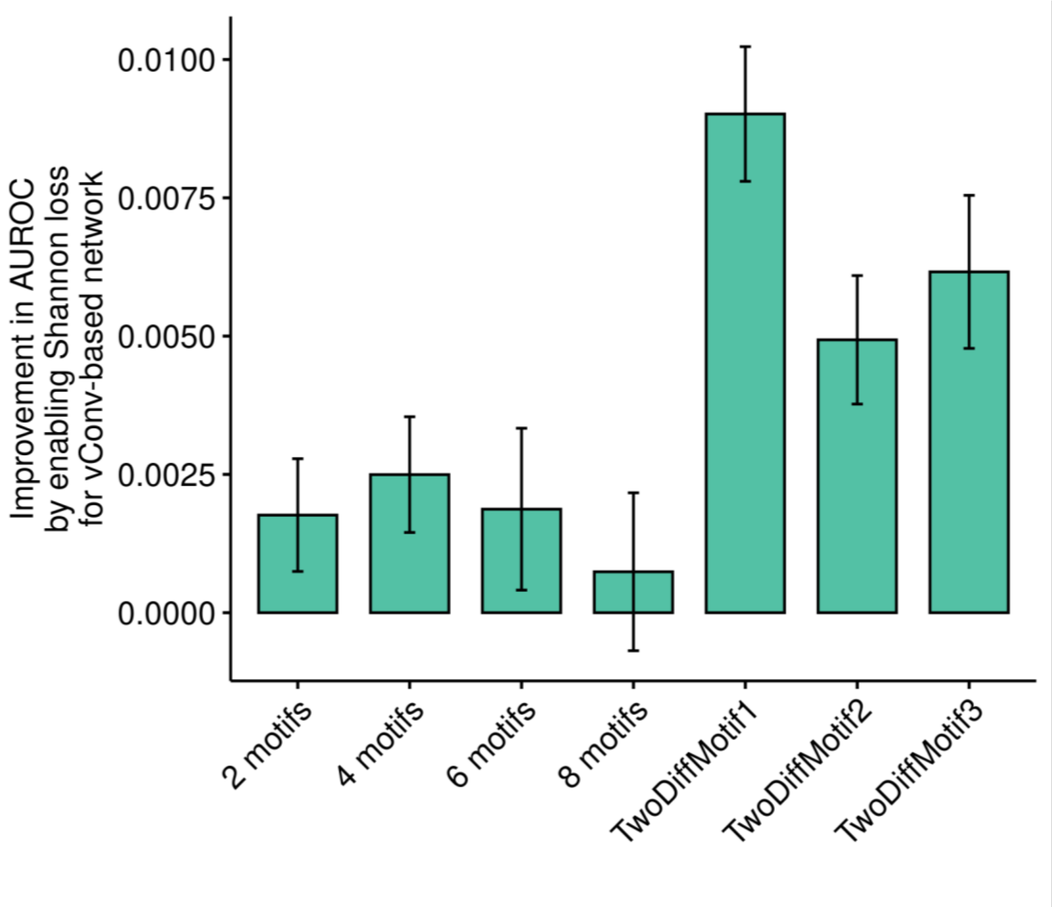


**Supplementary Fig. 11.** Comparison between vConv-based networks with or without Shannon loss across all hyper-parameter settings. The y-axis is the (per-hyper-parameter-setting) AUROC difference of the vConv-based network with Shannon loss minus that of the vConv-based network without Shannon loss.

A

$$Score(L_k|P_{ideal},\mathcal{M}) := E_K(P_{real}) = E_K(Prob(X_s > X_n|K,\mathcal{M}))$$

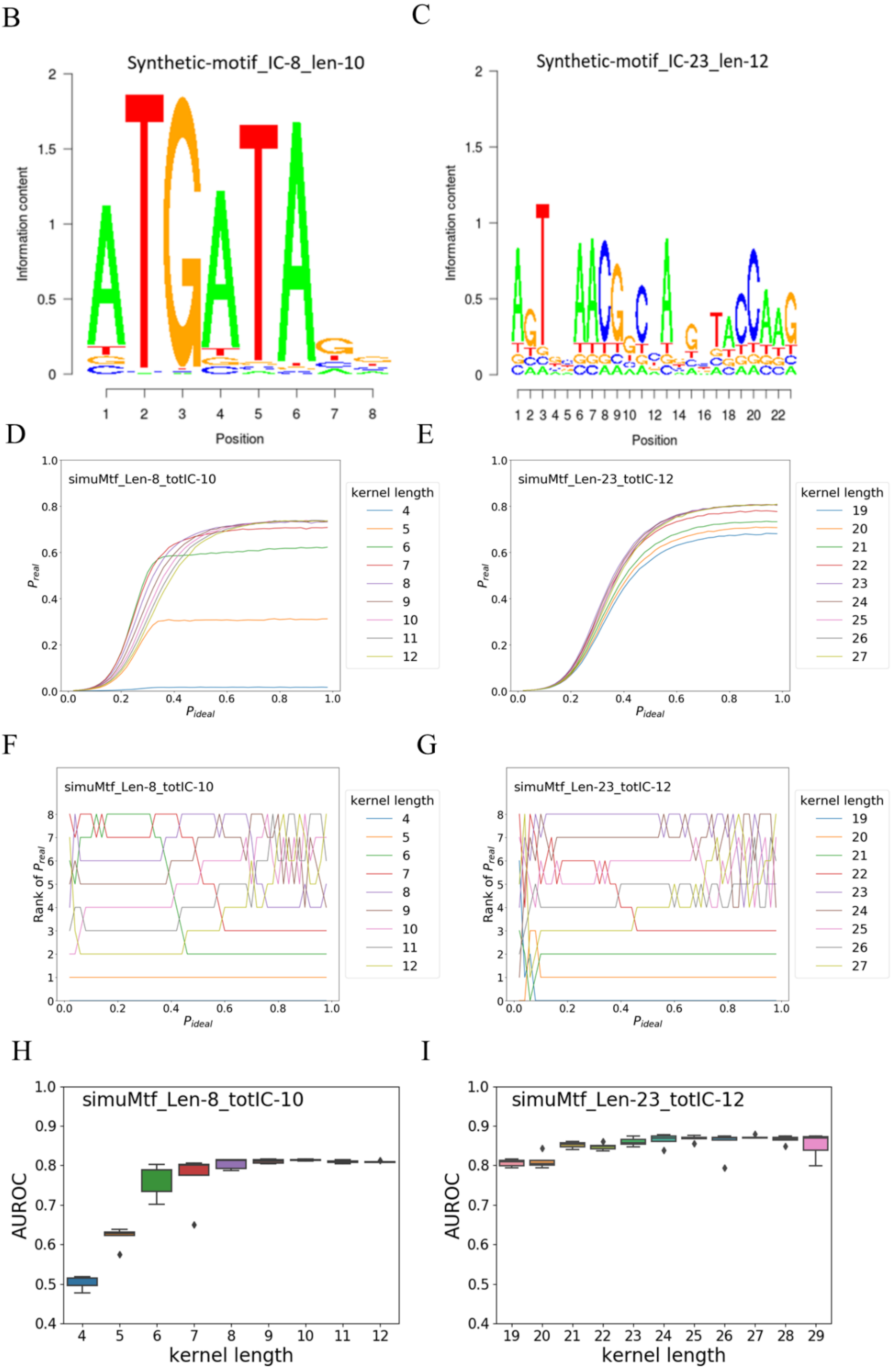

**Supplementary Fig. 12.** Theoretical modeling of the kernel in terms of the underlying real motif helps to evaluate the “goodness” of a particular kernel length for identifying the real motif. (A-C) We developed a scoring function (A) for calculating the kernel length and tested it on two example motifs, one of which (B) is shorter and more conserved than the other (C). (D-I) The kernel length did affect the model performance, although in a rather complicated way even in such a simplistic setting: for the first, shorter motif, the kernel length with the largest score depended on $P_{ideal}$ (D and F), while for the second, longer motif, the kernel length with the largest score was 23 (i.e., the motif length) for most $P_{ideal}$ values if we ignored the differences arising from numerical error (E and G). Similarly, we observed a complicated relationship between the kernel length and the average AUROC value of convolution-based networks in practice (H and I)
