## Supplementary material for "Identifying complex motifs in massive omics data with a variable-convolutional layer in deep neural network": Supplementary_tables_and_notes.pdf

| Hyper-parameter | Value range space for comparison in Fig. 3B | Value range space for comparison in Fig. 3A and Supplementary Fig. 7C |
|---|---|---|
| Kernel number | {128} | {128} |
| Kernel length | {24} | {10,17,24} |
| Random Seeds | 8 different seeds | 8 different seeds |

**Supplementary Table 1**. Hyper-parameter space for comparison between vConv-based and convolution-based networks on ENCODE datasets.

| Dataset | Std. for vConv-based networks | Std. for convolution-based networks | p-value |
|---|---|---|---|
| 2 motifs | 0.049 | 0.107 | <0.001 |
| 4 motifs | 0.053 | 0.097 | <0.001 |
| 6 motifs | 0.074 | 0.111 | <0.001 |
| 8 motifs | 0.078 | 0.096 | <0.001 |
| TwoDiffMotif1 | 0.051 | 0.132 | <0.001 |
| TwoDiffMotif2 | 0.054 | 0.117 | <0.001 |
| TwoDiffMotif3 | 0.063 | 0.149 | <0.001 |

**Supplementary Table 2**. The standard deviation (std) between vConv-based and convolution-based networks on each simulation dataset. The p-value is from the Levene's test with the null hypothesis that the variances of these two networks are equal.

| Dataset | Std. for vConv-based networks with MSL | Std. for vConv-based networks without MSL | p-value |
|---|---|---|---|
| 2 motifs | 0.049 | 0.05 | 0.112 |
| 4 motifs | 0.053 | 0.053 | 0.047 |
| 6 motifs | 0.074 | 0.073 | 0.793 |

| | | | |
|---|---|---|---|
| 8 motifs | 0.078 | 0.076 | 0.947 |
| TwoDiffMotif1 | 0.051 | 0.068 | <0.001 |
| TwoDiffMotif2 | 0.054 | 0.063 | <0.001 |
| TwoDiffMotif3 | 0.063 | 0.072 | <0.001 |

**Supplementary Table 3**. The standard deviation (std) between vConv-based networks with or without MSL on each dataset. The p-value is from the Levene's test with the null hypothesis that the variances of these two networks are equal.

**SUPPLEMENTARY NOTE 1: Kernel length affects the performance of convolution-based networks**

While such an effect has been suspected for a long time, no previous study has systematically investigated whether (and how) kernel length affects convolution-based networks' performance, especially when the underlying signals are of mixed lengths [1, 2]. In computer vision, researchers have empirically noticed that different kernel lengths in CNNs lead to differences in performance; for example, Han [3] reported that when a CNN is applied for facial action unit recognition (FAUR), changes in the CNN's kernel size will affect the performance of the model. Moreover, Han's results show that for FAUR, the optimal kernel size is different on different datasets, and there is no overall tendency for either a large or small kernel size to be preferred. Thus, the Inception model [4] tries to combine multiple kernels with different sizes for boosting global performance for various computation vision tasks [5, 6].

In addition to empirical assessments, we can further theoretically model the relationship between kernel length $(L_k)$ and model performance using a probability-based scoring function (Supplementary Fig. 11A; see "Mathematical treatment of the scoring function" below for the full mathematical treatment). Basically, for each combination of (1) a real $L_k$-by-4 motif $\mathcal{M}$, (2) the proportion ($P_{ideal}$) of the kernel contributed by this real motif (where the kernel is defined as $P_{ideal} * \mathcal{M} + (1 - P_{ideal}) * R$, with $R$ being a random matrix representing noise), and (3) $L_k$, this scoring function computes the expected probability across all possible kernels that the kernel's convolution with an arbitrary $\mathcal{M}$-containing sequence will take its maximal value at the position at which the motif is inserted. A high score for a certain $L_k$ indicates that kernels of this length can easily distinguish $\mathcal{M}$-containing sequences from other sequences, thus leading to good performance of the final convolution-based networks. The results of applying this scoring function to various cases (Supplementary Fig. 11B-G) clearly demonstrated that the scoring function helps to quantify the "goodness" of a particular $L_k$ for identifying a given motif under a particular $P_{ideal}$.

**Mathematical treatment of the scoring function**

Given (1) a real motif $\mathcal{M}$ (described by a PWM), (2) how much of the kernel is made up of the real motif ($P_{ideal}$, a scalar in the range of [0, 1]), and (3) the kernel length $L_k$ (note that it need not match the length of $\mathcal{M}$), we define the following scoring function:

$$\text{Score}(L_k|P_{ideal},\mathcal{M}) \coloneqq E_{\text{K}}(P_{real}) = E_{\text{K}}(Prob(X_s > X_n|\text{K},\mathcal{M}))$$

Where:

1. $\text{K}$ is a (trained) $L_k$-by-4 kernel matrix of length and is assumed to have the following form, where $\widetilde{\mathcal{M}}$ is a fixed-length version of $\mathcal{M}$ that is cut or padded to match the length of $\text{K}$ and $\text{Rand}$ is a random noise matrix of the same shape as $\text{K}$; the expectation ($E_{\text{K}}$) is thus taken over all possible $\text{K}$'s with different $\text{Rand}$'s.

$$\text{K} \coloneqq P_{ideal} * \widetilde{\mathcal{M}} + (1 - P_{ideal}) * \text{Rand}$$

2. $X_s$ is the value obtained by convolving $\text{K}$ with an arbitrary $\mathcal{M}$-containing sequence at the position where $\mathcal{M}$ is inserted.
3. $X_n$ is the maximum among all such values obtained by convolving $\text{K}$ with the same sequence mentioned above at positions other than the position where $\mathcal{M}$ is inserted.
4. $P_{real} \coloneqq Prob(X_s > X_n|\text{K},\mathcal{M})$ is the probability that the value obtained by convolving the kernel with an arbitrary $\mathcal{M}$-containing sequence will be the largest at the position where $\mathcal{M}$ is inserted.

The following steps are then applied to numerically compute the scoring function:

1. Determine $\widetilde{\mathcal{M}}$. If $\mathcal{M}$ is shorter than $\text{K}$ by $\text{d}$ nucleotides, then $\widetilde{\mathcal{M}}$ is padded with $\lfloor \text{d}/2 \rfloor$ and $\lceil \text{d}/2 \rceil$ [0.25, 0.25, 0.25, 0.25] columns on the left and right sides, respectively. If $\mathcal{M}$ and $\text{K}$ are of the same length, then $\widetilde{\mathcal{M}} = \mathcal{M}$. Otherwise, (i.e., if $\mathcal{M}$ is longer than $\text{K}$), $\widetilde{\mathcal{M}}$ is obtained by cropping $\mathcal{M}$ from both sides such that the sum of the information content in all positions of $\widetilde{\mathcal{M}}$ is the largest among all possible cropping outcomes.
2. Sample the random variables:
   a) The elements of $\text{Rand}$ are i.i.d. and are sampled from the uniform distribution $U(-0.25, 0.25)$.
   b) For each arbitrary $\mathcal{M}$-containing sequence, it is assumed to be one-hot encoded, its length is fixed to 1000, and the position of motif insertion, $\text{i} \in [1, 1000 - \text{motif length}]$, is sampled from the distribution $Prob(i) = \frac{1}{1000 - \text{motif length} + 1}$; then,

the motif part of this sequence is generated as specified by the PWM of $\mathcal{M}$, and all remaining positions in this sequence are i.i.d. and are sampled from the multinomial distribution [0.25, 0.25, 0.25, 0.25].

3. Calculate $X_s$ and $X_n$ via convolution. For the convolution operation, only positions where the kernel falls within the sequence are considered.
4. Obtain the final result. For a randomly generated pair of $\mathrm{K}$ and a random sequence, we can calculate their $X_s$ and $X_n$. Accordingly, we randomly sampled 100,000 kernels and corresponding sequences and then estimated $E_{\mathrm{K}}(P_{real})$ as $\frac{\text{number of times } X_s \text{ is larger than } X_n}{100{,}000}$.

To evaluate the relationship between the kernel length and the average AUROC value of a convolution-based networks (Supplementary Fig. 11F-G), we constructed a convolution-based network with a single layer of convolution with 32 kernels, a global-maxpooling layer, and a dense layer plus a sigmoid function. This network was trained 5 times with different random seeds for each kernel length, and its AUROC was then computed based on the validation dataset. Parameters were initialized using the Glorot uniform initializer [7], and we used AdaDelta [8] as the optimizer with default parameters. Early stopping on validation was used with patience=50.

For each motif, we simulated 6,000 sequences (with 3,000 positive and 3,000 negative) of length 1,000, picked 1200 as the test dataset randomly, and split the rest into 4,320 training and 480 validation sequences by setting “validation_split=0.1” in model.fit of Keras Model API (version 2.2.4)[9]; in this way, for each motif case, the same collection of training, validation, and test datasets was used for each model on same dataset. The ratio of counts of positive to negative sequences in the test dataset was within [0.8, 1.2] for both two motif cases. Each positive sequence is a random sequence with a “signal sequence” inserted at a random location, and each negative sequence is a fully random sequence. The “signal sequence” is a sequence fragment generated from one of the motifs in Supplementary Fig 11 B-C.
